# Evolution of Large Eyes in Stromboidea (Gastropoda): Impact of Photic Environment and Life History Traits

**DOI:** 10.1101/2024.04.09.588691

**Authors:** Alison R. Irwin, Nicholas W. Roberts, Ellen E. Strong, Yasunori Kano, Daniel I. Speiser, Elizabeth M. Harper, Suzanne T. Williams

## Abstract

Eyes within the marine gastropod superfamily Stromboidea range widely in size, from 0.2 to 2.3 mm, the latter being the largest known eyes of any gastropod. However, the evolutionary pressures underlying this variation remain unknown. Here, we use the wealth of material available in museum collections to explore the impact of ecological factors that affect light availability (ocean depth, turbidity and diel activity) on the evolution of eye size and structure within Stromboidea. We construct the most taxonomically extensive phylogeny of Stromboidea to date as a framework to investigate these relationships. Our results suggest that depth is a key light-limiting factor in stromboid eye evolution; here, increasing water depth is correlated with increasing aperture width relative to lens diameter (p = 0.043), and therefore an increasing investment in sensitivity in dim light environments. In the clade containing all large-eyed families (Strombidae, Rostellariidae and Seraphsidae), cathemeral species had wider eye apertures relative to lens sizes than diurnal species (p = 0.002), thereby prioritising sensitivity over resolution. These cathemeral species also had smaller body sizes than diurnal species (p = 0.002); this may suggest that animals with smaller shells are more vulnerable to shell-crushing predators, and therefore avoid the higher predation pressure experienced by animals active during the day.

Within the large-eyed stromboid clade, ancestral state reconstruction estimates that absolute eye size increased above 1 mm twice independently. Due to the high energetic investment associated with large eye sizes, this repeated increase in eye size suggests that higher performance vision is important in behavioural tasks within these families. In addition, we note that species with more robust escape responses tend to have larger eyes than those that do not: for example, strombids have 1.79 ± 0.31 times the eye size that would be expected from their body size (based on Stromboidea as a whole) and are known to display either a rapid escape response from an approaching predator or withdrawal into the shell. By contrast, the small-eyed xenophorids have 0.59 ± 0.30 times the expected eye size, and are only known to withdraw when threatened.

---

Eye size and structure vary enormously across the animal kingdom, with diverse adaptions driven by needs such as orienting through complex environments, finding suitable food and mates, and detecting potential predators. For animals relying on vision to complete these tasks, eyes must provide sufficient spatial resolution (ability to perceive detail) and sensitivity (ability to perceive contrast) (Land, 1981; Land and Nilsson 2012). However, adaptations to increase either resolution or sensitivity often come at the expense of the other (Warrant and McIntyre 1993). One solution to this trade-off is to increase the size of the eye, which, with all structural proportions kept constant, can increase both resolution and sensitivity (Land and Nilsson 2012). Larger eyes are associated with both longer focal lengths and wider apertures, the latter of which increases the amount of light entering the eye (Warrant 2004; Warrant and Locket 2004). However, a large eye requires a significant energetic investment during development and throughout life in encoding and processing visual information and continuous repair of light-induced photoreceptor damage (Laughlin et al. 1998; Meyer-Rochow 2001; Niven and Laughlin 2008). Therefore, large eye size is a characteristic of animals that rely heavily on vision, as the benefits of increased visual information must outweigh costs (Hiller-Adams and Case 1988; Moran et al. 2015; Nilsson et al. 2009, 2013).

Species that live in brighter environments typically have larger eyes, a trend well-documented in several groups (birds [Hall and Ross 2007]; insects [Somanathan et al. 2009]; reptiles [Hall 2008]; fish [Caves et al. 2017]; crustaceans [Beston et al. 2019]). In the marine realm, the availability of light is predominantly influenced by depth and water turbidity, with turbidity correlated with latitude (Warrant 2004). The attenuation of sunlight by seawater is a function of wavelength and due to processes of scattering and absorption, wavelengths longer than 600 nm and less than 400 nm are rapidly attenuated, even in the clearest of ocean waters. However, shorter wavelengths between 400 nm and 500 nm penetrate further resulting in a narrowband wavelength distribution. Therefore, at 200 m the intensity of light available for vision has declined by over three log units (Jerlov 1976). Animals inhabiting such low-light conditions often have larger eye sizes and aperture widths which increase sensitivity (Warrant et al. 2003; Warrant and Locket 2004), although the reverse trend is typically seen below 1,000 m depth where the only source of light is bioluminescence (Martin et al. 2007; Liu et al. 2012; Caves et al. 2017; Williams et al. 2022). Similarly, nocturnal activity is known to be associated with cellular-level adaptations such as longer rhabdoms, which also increase sensitivity (Blest and Land 1977; Narendra et al. 2017). Overall, however, there has been limited research on evolutionary relationships between light quantity and quality and eye size of marine invertebrates, and particularly in deep-sea taxa due to sampling difficulties (Hiller-Adams and Case 1988; Williams et al. 2022). Therefore, the accumulated material in museum collections, and their associated data, provide important opportunities for such studies, as has been demonstrated in other groups (Juarez et al. 2019; Thomas et al. 2020).

A striking example of evolution towards larger eye sizes is found within the marine gastropod superfamily Stromboidea. Mitogenome studies resolve two clades within the superfamily (Irwin et al. 2021; Machkour-M’Rabet et al. 2021; Li et al. 2022), one of which comprises three families, Strombidae, Seraphsidae and Rostellariidae, that are well known for their prominent and often colourful eyes on the ends of long, mobile eyestalks (for ease of reading, hereafter referred to as the ‘large-eyed clade’). By contrast, eyes of the remaining families in the second clade, Aporrhaidae, Struthiolariidae and Xenophoridae, are small and darkly pigmented and are situated at the base of cephalic tentacles (referred to as the ‘small-eyed clade’) (Woodward 1894; Simone 2005). These six families inhabit a diverse range of habitats that vary in depth and employ a variety of feeding and locomotory strategies (Supplementary Table S1 available on Dryad) (Berg 1975; Savazzi 1988; Ponder and de Keyzer 1998; Wells 1998). Behavioural and anatomical studies have demonstrated that large strombid eyes provide the highest spatial resolution of any mollusc except the Cephalopoda and Pterotracheoidea, and indeed among the finest resolution of any invertebrate (Seyer 1994; Irwin et al. 2022). This high-acuity vision supports early detection of predators in withdrawal responses (Irwin et al. 2022), and likely informs the unique escape response elicited in the presence of molluscivorous cone snails (Kohn and Waters 1966; Field 1977). However, it is not known when or how often large eye size evolved in the group.

In this study, we use specimens from museum collections to construct the most complete phylogeny for Stromboidea to date, comprising a third of all known species, and time-calibrated this tree using data from the rich stromboid fossil record. Within this phylogenetic framework, we used ancestral state reconstruction to explore the impact of depth, turbidity and diel activity on the evolution of eye size and other visual traits. The systematics of the group will be addressed in a future paper with increased taxon sampling (Irwin et al. in prep); however, the scope of the current study is suitable for identifying some of the key drivers underpinning the great morphological variation in stromboidean visual systems.

## Methods

### Sample Choice

Sampling for phylogenetic and morphological studies consisted of 72 specimens representing 56 species across Stromboidea, comprising 67% of currently recognised genera as listed in MolluscaBase (2023). These specimens were provided by colleagues or obtained from museum collections. See Supplementary Table S2 (available on Dryad) for collection data, GenBank numbers, and museum registration numbers for vouchers. The eyes of 40 species used in molecular studies were sufficiently well-preserved to obtain morphological data. For the remaining 16 species, molecular data were collected from one specimen, and morphological data from another specimen of the same species.

### Sequence Data

DNA was extracted from ethanol-preserved foot or mantle tissue according to the manufacturer’s instructions for the E.Z.N.A.® Mollusc DNA extraction kit (Omega Bio-tek). Fragments of the nuclear 28S rRNA gene (28S) and three mitochondrial genes, cytochrome oxidase subunit I (COI), 16S rRNA (16S), and 12S rRNA (12S), were amplified and sequenced following Williams and Ozawa (2006) (except that most 28S sequences were obtained with a novel forward primer, designed for Stromboidea; 5’– CAGTAACGGCGAGTGAAGC–3’). Sequencing was undertaken at the Natural History Museum, London (NHMUK), except for 13 COI sequences provided by the Muséum national d’Histoire naturelle (MNHN).

The dataset was comprised of sequence data for all four genes, where available (98% gene coverage; Supplementary Table S2). Alignment was performed via PASTA (Mirarab et al. 2014) and removal of ambiguously aligned regions by Gblocks (Castresana 2000) following Irwin et al. (2021); alignment of COI was unambiguous. The COI dataset of 658 bp included 268 variable sites, of which 257 were phylogenetically informative. After removal of ambiguous blocks, the alignment for 12S was 542 bp (85% of 640 bp), with 274 variable sites and 227 phylogenetically informative sites; 497 bp for 16S (90% of 554 bp), with178 variable sites and 146 phylogenetically informative sites; and 1363 bp for 28S (84% of 1630 bp), with 337 variable sites and 220 phylogenetically informative sites.

### Morphological Trait Data

#### Data collection: micro-computed tomography and histology

Eye trait data were collected from adult snails using rapid micro-computed tomography (µCT; 50 species) or histology (6 species), with one eye used for each species (Supplementary Table S2; see Supplementary Table S3 for measurements available on Dryad). Specimens were confirmed as adults by a thickening of the terminal varix (shell outer lip). The best-preserved, intact eye was selected, except where ethanol-preserved molecular specimens had eyes removed upon collection and preserved separately in glutaraldehyde; this glutaraldehyde-preserved material was used preferentially for histology (Supplementary Table S2). For consistency, the left eye was used for morphological examination when possible, although measurements taken from images via Fiji v. 1.52p (Schindelin et al. 2012) show no significant difference between left and right eyes for mean whole eye diameter (i.e., eye diameter including eyestalk tissue; Supplementary Fig. S1 available on Dryad), according to a Wilcoxon test performed in R v.4.0.3 (R Core Team 2022) (W = 1460.5, n = 54, p = 0.990; see Supplementary Table S4 (available on Dryad) for specimens included in analysis). Measurements of morphological traits, as described below, were subsequently log-transformed (log base 10) for all downstream analyses.

Methods for µCT-scanning followed Sumner-Rooney et al. (2019), with minor modifications in PTA concentrations and staining time, as well as agarose concentrations. Samples were stained in 1.5% phosphotungstic acid in ethanol for 7–19 days, then placed inside 0.2 ml PCR tubes and immobilised using 1% agarose topped up with ethanol. Each tube was placed within a 1,000 µl pipette tip fixed to a mount. Regions of interest were imaged with a Carl Zeiss Xradia Versa 520 (Pleasanton, United States) at NHMUK. Two scanning methods were used: first, higher resolution scans were taken of eyes from 1–2 representatives from each family for reconstructive purposes, included in this study as additions to the morphological dataset (Supplementary Table S2). These were conducted at 40 kV and 75–76 mA, taking 3,201 projections with an exposure time of 8 s (voxel size 2.5– 3.8 μm, identical in each plane e.g. 2.5 x 2.5 x 2.5 μm^3^). Rapid scans were used for remaining specimens, conducted at 40–70 kV and 75–86 mA, taking 1,001–1,601 projections with an exposure time of 2–5 s (voxel size 3.3–7.5 μm). Scans were viewed (and some manually segmented for comparative work) in Volume Graphics VGStudio Max v. 2.2, with morphological measurements taken as described below.

For histology, samples stored in 2.5% glutaraldehyde in 0.1M phosphate-buffered solution (Supplementary Table S2) were dehydrated in a graded ethanol series (20 minutes in 30, 50, 70, 80, 95 and 100% ethanol). Ethanol-preserved samples (Supplementary Table S2) only required dehydration for 20 minutes in 95 and 100% ethanol. All samples were subsequently dehydrated in 100% acetone (2 x 20 minutes), transferred to propylene oxide (3 x 20 minutes), then to 1:1 propylene oxide and Spurr’s resin (hardness A) overnight (Spurr 1969). This was replaced by 100% Spurr’s resin (4–24 hours, depending on the sample size), then the sample was embedded for 48 hours at 70° C. Samples were serially sectioned with a diamond knife (1 μm sections; DiATOME® Histo Jumbo 8 mm) on an automated microtome (Sorvall^TM^ RMC MT6000), and stained using Richardson’s solution (Ruthensteiner 2008; modified from Richardson et al. 1960). Sections were imaged using an Olympus BX63 light microscope (NHMUK).

#### Morphological trait measurements

A trait analysis of this broad taxonomic scale can only be achieved using museum collections, so measurement methods were designed to mitigate differences in preservation (Supplementary Table S2). Mitigating steps are described below for each trait where relevant, however, as most ethanol-preserved specimens (64%) and all specimens preserved in glutaraldehyde used in morphological analyses were collected in the past 10 years, we consider variation in specimen condition to be a minor issue when analysing overall trends in the data. Moreover, other studies investigating trends in eye evolution using museum specimens found tissue shrinkage due to preservation to be a minor concern (Thomas et al. 2020).

Absolute eye diameter was measured as the maximum width of the eye bulb, including all visual structures within the capsule (see Supplementary Figure S1 for illustrations of measurements). Preservation differences caused shrinkage of the vitreous body in some specimens, resulting in part of the eye bulb pulling away from the eyestalk; when this occurred, the empty cavity was included in eye width measurements. Relative eye diameter was calculated as the ratio of eye diameter to a body size proxy. Body size was estimated using the shell body-whorl width, measured ventrally at the widest point of the last whorl (excluding the flared outer lip, avoiding calluses or spines when possible, and measured perpendicular to the growth lines of the shell) (Supplementary Fig. S1). For xenophorids, which have a trochiform shell shape as opposed to the fusiform shell shape of other stromboids (Simone 2005), the widest point of the last whorl was measured from the lateral view, parallel to the flat base of the shell, and avoiding agglutinated objects (Supplementary Fig. S1).

Morphological data so far have shown (Gillary and Gillary 1979; Seyer 1994; Blumer 1996; this study) that stromboids possess a spherical lens. However, preservation seems to cause lenses to distort in unpredictable ways; therefore, lens diameter was calculated from the lens volume, taken via VGStudio Max assuming a sphere (Supplementary Fig. S1).

Eye aperture diameter was measured in Fiji from images taken directly above the eye by a Zeiss Axio Zoom microscope (taken along axis of whole eye diameter; Supplementary Fig. S1). Initial comparisons of aperture measurements between these images and histological sections of the same specimens confirmed their accuracy. To investigate relative aperture size, the ratio of lens diameter to aperture width was calculated.

Rhabdom length was averaged from 18 measurements of the rhabdom layer (individual cells were not distinguishable) from a single eye, taken in the sagittal and frontal sections at evenly spaced intervals along the basal third of the eye bulb, where preservation was most consistent (Supplementary Fig. S1). Measurements were not taken at points where the retina had pulled away from the eyestalk during preservation. Note that although screening pigment was not visible in μCT scans, the rhabdom layer was distinguishable from the nuclear layer due to a difference in contrast. To compare investment in resolution versus sensitivity, relative rhabdom length was calculated as the ratio of lens diameter to rhabdom length.

### Habitat Trait Data

Sources of additional data (e.g. museum databases) are listed below with respect to each habitat trait. For the five putative cryptic species included in this study, collection data were only used if specimens had molecular data available, so that habitat traits could be correctly assigned to each cryptic species. Exceptions were made for *Canarium mutabile* B and *Canarium wilsonorum* A, which were identified as allopatric to their respective sister taxa (based on available data; Irwin et al. in prep); for these species, data from specimens collected in the same regions as molecular specimens were also used.

#### Depth

Maximum and minimum depth data were sourced from the literature (see Supplementary Table S3). Additional data from museum collection databases were used to supplement depth range data, in addition to specimens in private collections featured on http://www.stromboidea.de/ (Wieneke et al. 2022); see Supplementary Table S3 for details. Depth uncertainty associated with trawl sampling was accounted for by taking the shallowest of the deepest values as the maximum depth, and the deepest of the shallowest values as the minimum depth (following Bouchet et al. 2008).

#### Diel activity pattern

Species were scored as cathemeral (1) when at least one specimen was recorded as hand-collected at night (assumed to be active), or when species identity could be verified in dive photos taken at night of active animals assumed to be foraging, or when literature reported animals as active at night (see Supplementary Material 6 for justifications). When all the above were lacking, species were assumed to be active during the day only, or else significantly less active at night, and scored as (0). Species inhabiting maximum depths of ≥ 200 m where light levels are lower were scored as (1) by default. Due to morphological and geographical similarity among seraphsid cryptic species, the records found for *Terebellum* were used to score both included species as (1) (Supplementary Table S5 available on Dryad).

#### Turbidity

The diffuse attenuation coefficient at 490 nm (Kd490) is a measure of the rate at which visible light in the blue to green region of the spectrum is attenuated with depth and is used as an indicator of turbidity. A Moderate Resolution Imaging Spectroradiometer (MODIS) Aqua mission composite map of Kd490 data from 01–01–2003 to 01–01–2022 at 4 km resolution (OBPG 2022) was overlain onto a global map via QGIS v. 3.22 (QGIS Development Team 2022). Specimen locality data were compiled from museum collections databases for each species and mapped onto the Kd490 data. Where at least one specimen was collected within an area of high turbidity, the species was scored as (1); where no locality points fell in areas of high turbidity, species were scored as (0). Based on initial observations, high turbidity is here defined as Kd490≥0.5 m^-1^, with lower turbidity as <0.5 m^-1^. This cut-off value is the midpoint of the ‘slightly murky’ mesotrophic and oligotrophic freshwater and coastal habitats (0.1<K<1.0) defined by Caves et al. (2017).

#### Phylogenetic Analysis and Simultaneous Ancestral State Reconstruction

A species-level phylogeny was produced using Bayesian Inference as implemented in BEAST v.1.10.4 (Drummond and Rambaut 2007). The dataset was partitioned by locus, with each gene allowed to evolve at a different rate. The best nucleotide substitution models were determined by ModelTest-NG (Darriba et al. 2020) using the Akaike Information Criterion (AIC) to be GTR+I+G for 28S, and HKY+I+G for 16S, and TrN93+I+G for COI and 12S. The tree was calibrated using four fossil dates listed in Table 1 (see Irwin et al. (in prep) for justifications of dates used). We used a Yule speciation prior with a *BEAST (v.2.6.6; Bouckaert et al. 2019) tree produced via the Species Tree Ancestral Reconstruction method from (Irwin et al. in prep) as the starting tree, trimmed via TreeGraph2 v. 2.15.0-887 beta (Stöver et al. 2010) to include only species used in this analysis. The BEAST analysis was run for 150,000,000 generations, sampling every 3,000 generations. Log files were examined for convergence via Tracer, and effective sample size (ESS) values >200 indicated adequate sampling. The final species tree produced via *TreeAnnotator* (BEAST package) was a maximum clade credibility tree with median node heights based on 45,000 trees, after a 10% burnin. Support for nodes was determined using posterior probabilities (PP; calculated by BEAST).

**Table 1.**
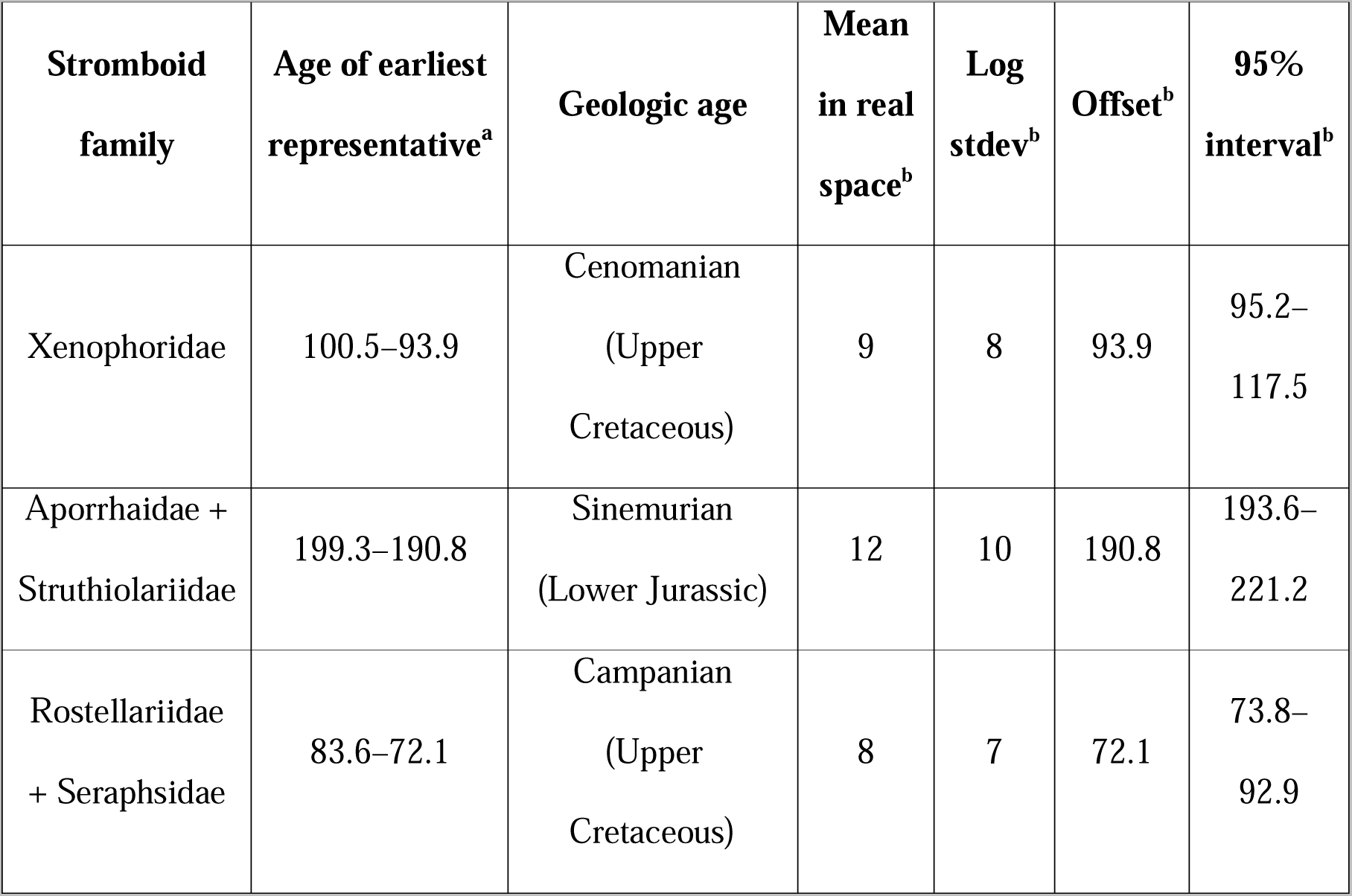
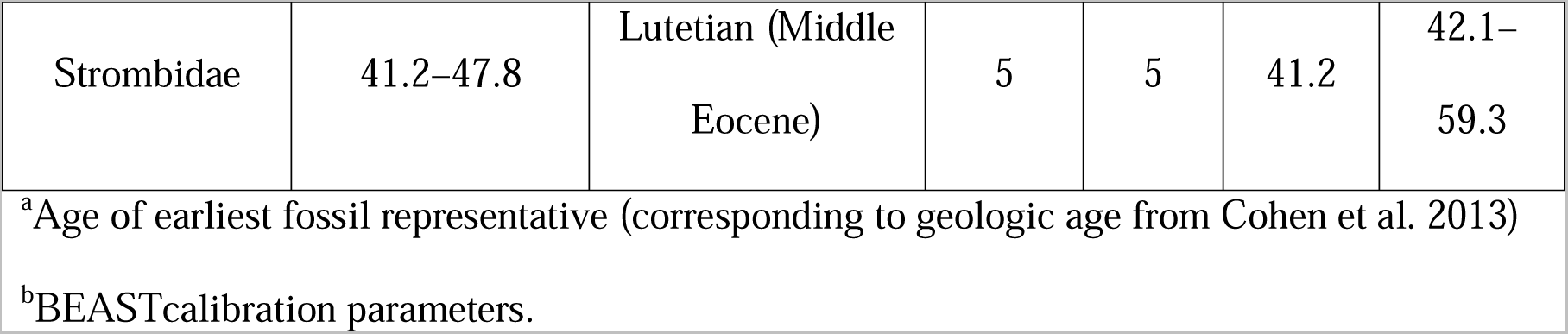
Fossil Calibrations Used in BEAST Analyses.

We inferred a continuous phylogeographic diffusion model for all continuous trait data using a Brownian random walk model (Gill et al. 2017). Discrete trait data (diel activity and turbidity) were analysed using a symmetric substitution model with equal forward and reverse rates, in association with the Bayesian Stochastic Search Variable Selection (BSSVS) procedure. Ancestral state reconstruction was performed for all traits, and state change counts were reconstructed for discrete data using Markov Jumps (Minin and Suchard 2008).

### Evolution of Traits: Phylogenetic Signal

We tested for phylogenetic signal for discrete traits (turbidity and diel activity) using the statistic *D* (Fritz and Purvis 2010), as implemented in R package *caper* (v. 0.5.2; Orme et al. 2012). This observed value of *D* can be tested for significant departure from two models (random association and the clumping expected under a Brownian evolution threshold model), and indicates whether a trait is liable to change or highly conserved. We examined continuous traits using the R package *phylosignal* (Keck et al. 2016), which computes five indices for general measurements of phylogenetic signal in continuous data: Moran’s *I* (Moran 1948, 1950), Abouheif’s *C_mean_* (Abouheif 1999), Blomberg’s *K* and *K\** (Blomberg et al. 2003) and Pagel’s λ (Pagel 1999). Each index is tested against the null hypothesis, H_0_, that trait values are randomly distributed across the phylogeny) (Keck et al. 2016). We also calculated the Local Indicators of Phylogenetic Association (Local Moran’s *I*) via function *lipaMoran* for each tip of the phylogeny for continuous traits, which performs a nonparametric test by randomisation for each value of *I* (Keck et al. 2016).

### Evolution of Traits: Correlation between Traits

#### Stromboidea

We evaluated continuous traits (absolute and relative eye diameter, absolute and relative aperture width, lens diameter, absolute and relative rhabdom length) as phylogenetically non-independent variables of discrete traits (turbidity and diel activity) using the *caper* function *brunch*. The *caper* function *crunch* was used to evaluate these morphological traits as variables of continuous maximum depth. Significance of the phylogenetically independent contrast (PIC) values generated in *brunch* and *crunch* was determined using a one-tailed *t*-test and *F*-statistic to investigate whether the mean of the independent contrasts was significantly different from zero.

Boxplots of minimum and maximum depth ranges were plotted for each genus or clade using R package *ggplot2* (Wickham et al. 2016). Kruskal-Wallis ranked sum tests were used to test for significant differences in minimum and maximum depth among genera, and absolute and relative eye diameter among genera. Ranked sum tests were also used to test for differences among genera in eye investment (residuals from a phylogenetic generalized least-squares (PGLS) regression for eye size versus body size, following Thomas et al. 2020). Tukey multiple pairwise-comparisons were used to determine which pairs of genera differed in these factors. Wilcoxon tests were used to compare the means of depth ranges, eye investment, absolute and relative eye diameter, between large-eyed ((Rostellariidae + Seraphsidae) + Strombidae) and small-eyed ((Aporrhaidae + Struthiolariidae) + Xenophoridae) clades. Chi-square tests were used to test for significant associations between diel activity turbidity with either group (i.e., large-eyed or small-eyed clade) or genus.

We used ANCOVA models to test whether taxon group (large- or small-eyed family) had a significant impact on (1) absolute eye diameter after controlling for body size (i.e., eye diameter ∼ body size + group); (2) relative eye diameter after controlling for body size; (3) rhabdom length after controlling for lens diameter. For the relationship between aperture width and lens diameter, the ANCOVA assumption of homogeneity of coefficients was not met (F = 2.231, p = 0.030); thus, a multiple regression model was used to test whether the regression slope was similar in large- and small-eyed groups (aperture ∼ lens width * group).

#### Large-eyed clade ((Rostellariidae + Seraphsidae) + Strombidae)

Student’s or Welch’s t-tests were used to compare the means of morphological traits between large-eyed taxa observed active during both the day and night, and those observed active only during the day. Only species living in brightly lit habitats were compared, so that any remaining morphological variation could be attributed to differences in diel activity. Species present in at least one area of high turbidity (*Strombus alatus* and *Thetystrombus latus*) were excluded, as were deeper-water species *Rimellopsis laurenti* and *Rostellariella delicatula* (depth ranges 59–280 m and 100–575 m, respectively).

## Results

### Phylogenetic Analyses

The BEAST analysis is largely consistent with mitogenome studies (Irwin et al. 2021; Machkour-M’Rabet et al. 2021; Li et al. 2022), except for the position of *Varicospira* which is unresolved in this study (Fig. 1). Likewise, this study recovered two clades: a large-eyed clade ((Rostellariidae + Seraphsidae) + Strombidae), and a small-eyed clade ((Aporrhaidae + Struthiolariidae) + Xenophoridae) (Fig. 1), as in previous studies (Irwin et al. 2021; Machkour-M’Rabet et al. 2021; Li et al. 2022). Log files examined via Tracer revealed that 12S pInv and alpha traces continually switched between two states. Changing site model did not impact this; therefore, the analysis was run until all ESS values reached > 200.

**Figure 1.**
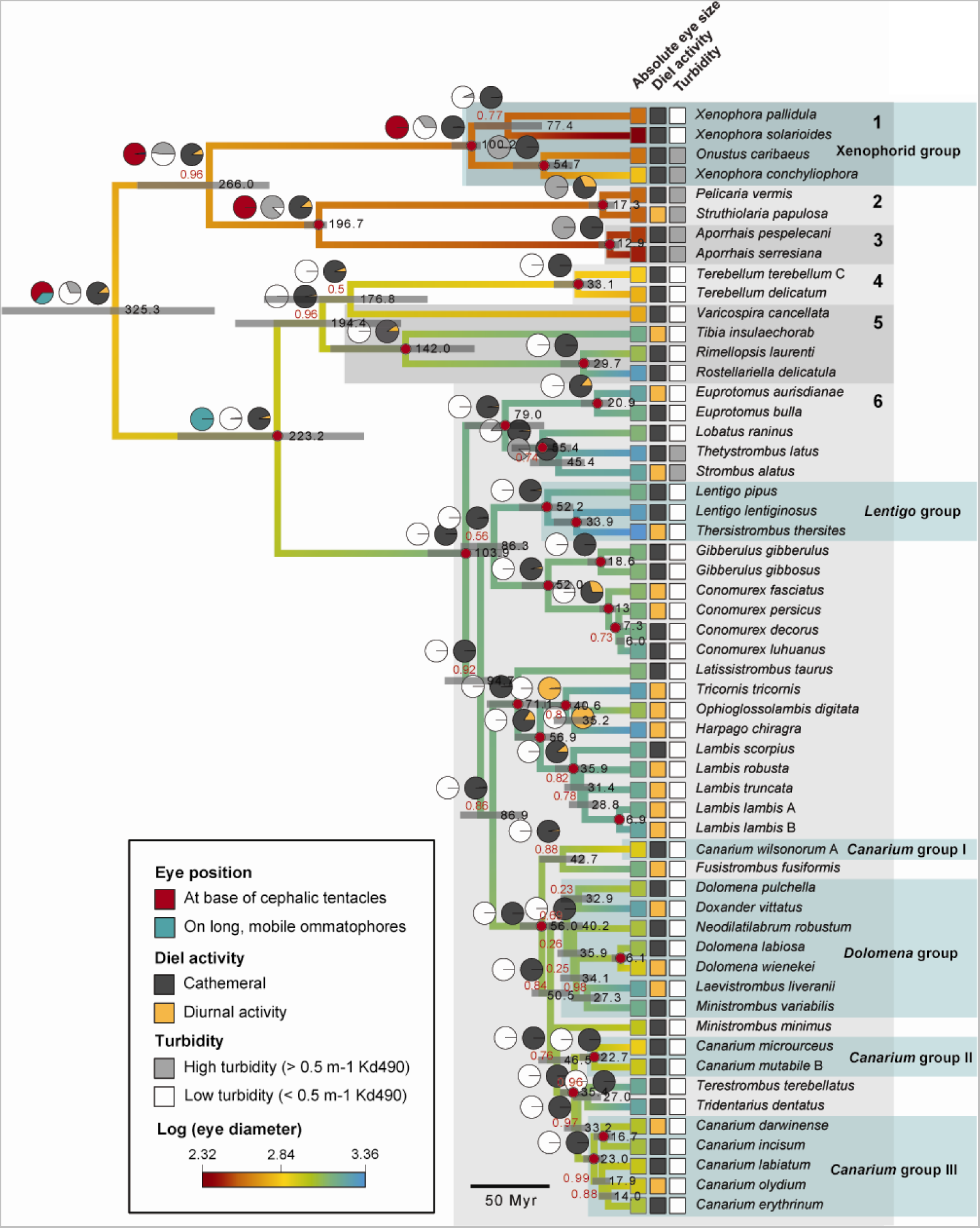
Bayesian analysis of 28S, COI, 16S and 12S sequences from 56 stromboid species as implemented in BEAST. Branch colours reflect log-transformed absolute eye diameter as reconstructed by ancestral state reconstruction (see methods for trait definition); red branch colour represents small eye diameter values and blue, larger values (see key). Numbers above branches (red) are PP; red circles at nodes indicate PP = 1.0. Numbers at nodes (black) are node ages; grey bars at nodes are 95% highest posterior density intervals (HPD) for node ages. Turbidity, diel activity (ancestral state reconstruction of discrete traits, except placement of eyes), and eye diameter are indicated for each species at tips by box colour (discrete traits: darker colour indicates trait presence, lighter colour indicates trait absence; see key). Pie diagrams indicating likely ancestral state reconstruction for discrete traits are shown for selected clades, corresponding to PP support for each character state; placement of eyes only shown for deep nodes at small- and large-eyed clades. Differently shaded grey boxes delimit families: 1, Struthiolariidae; 2, Aporrhaidae; 3, Xenophoridae; 4, Seraphsidae; 5, Rostellariidae; 6, Strombidae. For the purposes of generic- or clade-level statistical analyses, generic boundaries were adjusted according to the BEAST analysis results to prevent treating non-monophyletic genera as a clade; note use of the terms ‘*xenophorid* group’, ‘*Lentigo* group’, ‘*Canarium* groups I-III’ and *‘Dolomena* group’ for ease of discussion.

Analyses suggest the ancestral root node state for eye size in Stromboidea did not occupy either end of the extreme range of eye sizes represented by the extant stromboids (Table 2, Fig. 1). Rather, this was an animal with an intermediate absolute eye diameter (similar in size to the eyes of extant *Varicospira*, *Terebellum* and *Xenophora conchyliophora*), inhabiting a relatively shallow minimum depth and intermediate maximum depth (medians 4–155 m) (Table 2; Figs 1, 2). The stromboid ancestral state also had low support for a habitat of high turbidity (PP = 0.69), and moderate support for activity both at night and during the day (PP = 0.89) (Fig. 1).

**Figure 2.**
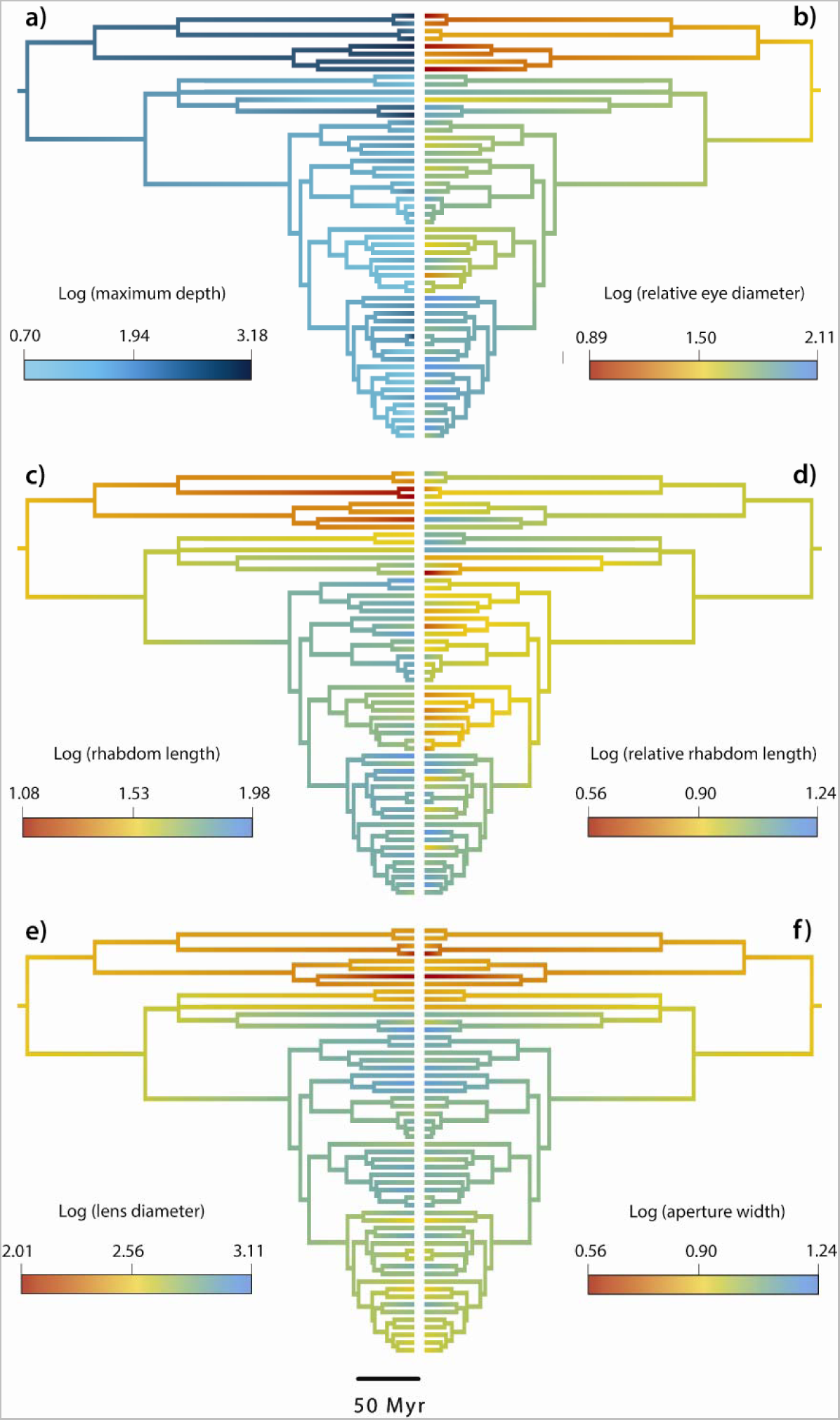
Further ancestral state reconstruction from Bayesian analysis as implemented in BEAST (see Fig. 1). Branch colours reflect log-transformed traits (see methods for full explanations of each trait): a, maximum depth; b, relative eye diameter; c, absolute rhabdom length; d, relative rhabdom length; e, lens diameter; f, absolute aperture width. For maximum depth, darker hues indicate increasing depth; for all morphological traits, red branch colour represents small values and blue, larger values.

**Table 2.** Ancestral Root Node State for Stromboidea (Median and Confidence Intervals Given).

|  | Median |  | 95% HPD |  |
| --- | --- | --- | --- | --- |
| Trait | Log units | Equivalent actual size<sup>a</sup> | Log units | Equivalent actual size<sup>a</sup> |
| Absolute eye diameter | 2.78 | 0.60 mm | 2.55–3.01 | 0.35–1.02 mm |
| Relative eye | 1.49 | NA | 1.23–1.74 | NA |

| diameter |  |  |  |  |
| --- | --- | --- | --- | --- |
| **Lens width** | 2.52 | 331.13 $\mu\text{m}$ | 2.30–2.75 | 199.53–562.34 $\mu\text{m}$ |
| **Absolute aperture width** | 2.48 | 302.00 $\mu\text{m}$ | 2.27–2.70 | 186.21–501.19 $\mu\text{m}$ |
| **Absolute rhabdom length** | 1.48 | 30.20 $\mu\text{m}$ | 1.34–1.64 | 21.88–43.65 $\mu\text{m}$ |
| **Minimum depth** | 0.63 | 4.27 m | 0.02–1.27 | 1.05–18.62 m |
| **Maximum depth** | 2.19 | 154.88 m | 1.72–2.66 | 52.48–457.09 m |

<sup>a</sup>Both log units and equivalent normal values are given for all states except relative eye diameter, where the normal value is a ratio of absolute eye diameter to body size.

Results suggest that the ancestral root node state of the large-eyed clade was an animal with a relatively large eye diameter (median 2.95 log units (= 0.89 mm), 95% HPD: 2.77– 3.12 log units (= 0.58–1.32 mm)). Remarkably, eye size increased further above a median of 1.00 mm in two separate clades within this large-eyed group: Strombidae (median 3.14 log units (= 1.38 mm), 95% HPD: 3.03–3.24 log units (= 1.07–1.74 mm)), and the rostellariid clade containing (*Rimellopsis* + *Rostellariella*) *+ Tibia* (median 3.04 log units (= 1.11 mm), 95% HPD: 2.86–3.22 log units (= 0.72–1.66 mm)) (Fig. 1). Eye size is also suggested to have decreased early in the evolution of the small-eyed clade, to a median of 0.49 mm at the root node of the clade around 265.96 Ma (Fig. 1). It is important to note that the time-calibrated analysis suggests an older radiation for Strombidae (103.9 Ma (95% HPD: 80.9–127.9 Ma) and Rostellariidae + Seraphsidae (194.4 Ma (95% HPD: 145.0–249.1 Ma) than is known from the fossil record (Fig. 1); the oldest fossil representatives for Strombidae and Rostellariidae date from the Middle Eocene and Campanian, respectively [*Stromboconus suessi* (Bayan 1870; Wieneke et al. 2022) and *Calyptraphorus itamaracensis* (Muñiz 1993)].

### Evolution of Traits: Phylogenetic Signal

Analyses suggested significant phylogenetic signal with respect to diel activity (*D* = 0.887, P_1_ = 0.299, P_0_ = 0.001; n_N_ = 36, n_D_ = 20); here, phylogenetic signal is defined as per Blomberg et al. (2003), i.e., that closely-related taxa are more likely to share a common trait than species selected at random from the tree. By contrast, results for turbidity indicated a random distribution across the tree (*D* = −0.829, P_1_ <0.001, P_0_ = 0.943; n_H_ = 8, n_L_ = 48). Here: n_H_ and n_L,_ number of species at high and low turbidity, respectively; n_N_ and n_D,_ number of species observed active both at night and during the day, or during the day only, respectively; P_1,_ probability of *D* score resulting from no phylogenetic structure; P_0,_ probability of *D* score resulting from Brownian phylogenetic structure.

The following continuous trait variables suggested a significant phylogenetic signal for all five indices tested: absolute and relative eye diameter, absolute aperture width, lens diameter and absolute rhabdom length (Table 3). For other traits, only the following indices showed significant signal: maximum depth, λ and *C_mean_*; minimum depth, λ; body size, *K* and *K\**; relative rhabdom length, *C_mean_* and *I* (Table 3). The LIPA analysis suggested significant autocorrelation between phylogeny and lens diameter, absolute aperture width and absolute eye diameter in the small-eyed clade (Supplementary Fig. S2 available on Dryad). Significant autocorrelation between phylogeny and relative eye diameter was also present in the clade containing the *Dolomena* group, *Canarium* s.l., *Tridentarius* and *Ministrombus* (but not *Terestrombus* or *Fusiformis*; Supplementary Fig. S2). In the clade containing *Latissistrombus + (((Tricornis + (Ophioglossolambis + Harpago)) + Lambis),* significant autocorrelation was present between all traits except relative eye diameter and absolute rhabdom length for many taxa in the group (Supplementary Fig. S2).

**Table 3.** Continuous Log-transformed Traits Examined Using *phylosignal*, Which Computes Five Indices for General Measurements of Phylogenetic Signal in Continuous Data.

|  | **Indices<sup>a</sup>** |  |  |  |  |
| --- | --- | --- | --- | --- | --- |
| **Trait** | $C_{mean}$ | $I$ | $K$ | $K^*$ | $\lambda$ |
| **Maximum depth** | 0.175* | -0.007 | 0.133 | 0.158 | 0.365** |
| **Minimum depth** | 0.094 | -0.004 | 0.140 | 0.164 | 0.344*** |
| **Body size** | 0.138 | 0.013 | 0.184*** | 0.275*** | 0.465 |
| **Absolute eye diameter** | 0.525*** | 0.099** | 0.519*** | 0.486*** | 0.789*** |
| **Lens diameter** | 0.481*** | 0.095*** | 0.433*** | 0.412*** | 0.715*** |
| **Absolute aperture width** | 0.444*** | 0.092** | 0.424*** | 0.393** | 0.667*** |
| **Absolute rhabdom length** | 0.496*** | 0.101** | 0.432*** | 0.389** | 0.6*** |
| **Relative eye diameter** | 0.393*** | 0.077** | 0.347*** | 0.386*** | 0.654*** |
| **Relative** | -0.018 | -0.021 | 0.147 | 0.215 | 0.00007 |
| **aperture**<br>**width** |  |  |  |  |  |
| **Relative**<br>**rhabdom**<br>**length** | 0.231** | 0.041** | 0.164 | 0.245 | 0.516 |

<sup>a</sup>Table contains values of statistics and associated p values for each tested trait and
method; asterisks mark significant p values (\*\*\*, $p \leq 0.001$ ; \*\*, $p \leq 0.01$ ; \*, $p \leq 0.05$ ).

### Evolution of Traits: Correlation between Visual Traits and Light Environment Stromboidea

In contrast to the small-eyed clade, large-eyed stromboid taxa generally inhabit environments with a high light availability, characterised by shallow maximum depths [mean 45.0 ± sd 32.5 m (s.e.m 4.8 m, n = 45, excluding deep-water taxa *Rimellopsis, Rostellariella* and *Dolomena labiosa*)] (Figs 2a, 3; Supplementary Table S3). Results suggested a significant difference between the means of maximum depth ranges for small- and large-eyed clades (*t*-test, *t* = −6.853, df = 54, p<0.001); among genera, these differences were narrowly not significant (Kruskal-Wallis, *H* = 41.127, df = 28, p = 0.052). The range and median of maximum depths for genera show that only some species within *Aporrhais*, the *Dolomena* group*, Rostellariella, Rimellopsis, Pelicaria,* and Xenophoridae extend below 200 m, with the latter two present below the dysphotic zone (200–1,000 m) (Fig. 3). In Tukey multiple pairwise-comparisons between genera/clades, the only significantly different maximum depths were between Xenophoridae and the following groups: *Lambis* (p = 0.003)*, Conomurex* p = 0.006)*, Canarium* group II (p = 0.04) and *Canarium* group III (p = 0.002). No significant associations were found with respect to minimum depth.

**Figure 3.**
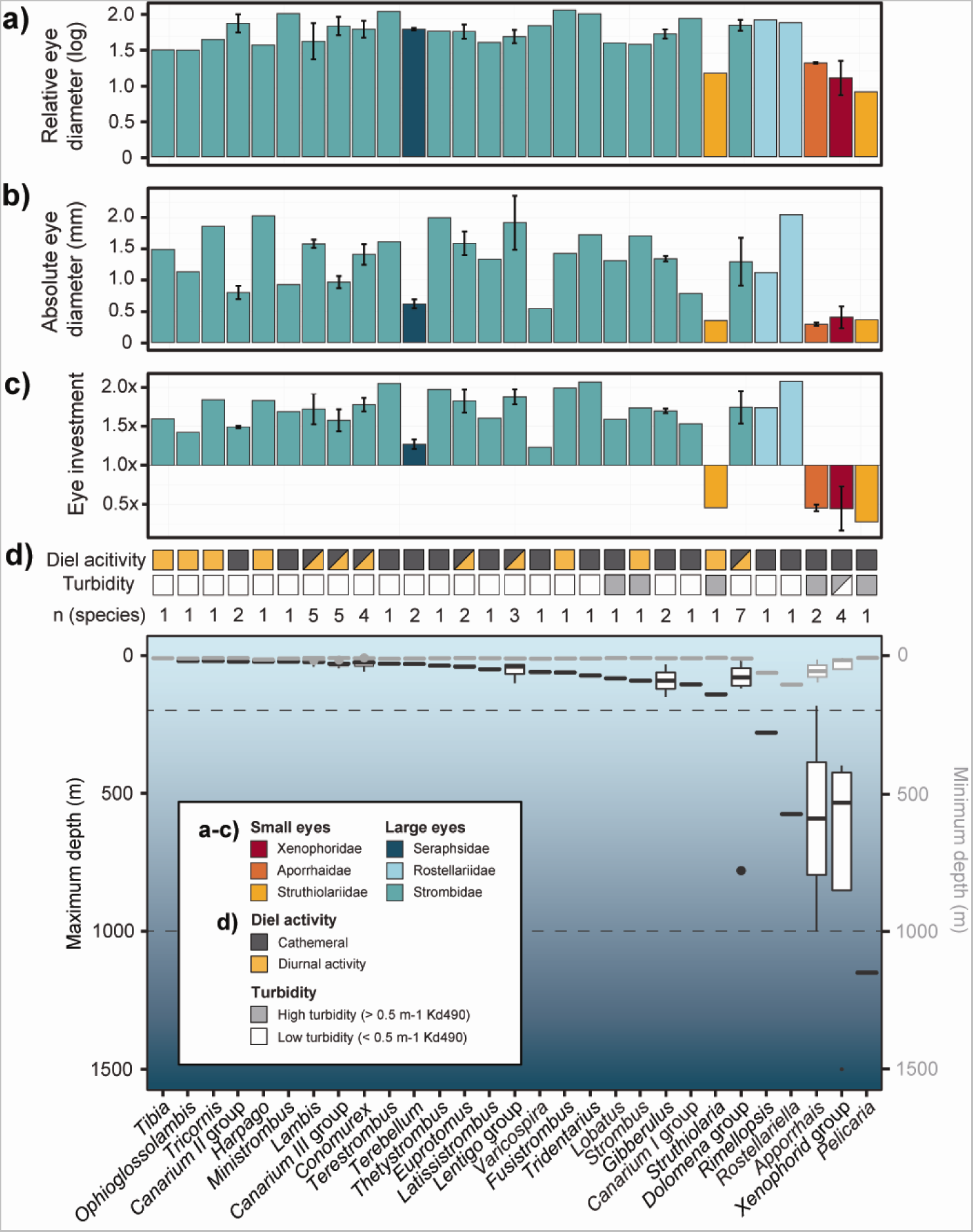
Variation in eye size and eye investment by light environment. Bar plots (a-c) (coloured by family; see key) show: a, log-transformed relative eye diameter; b, log-transformed absolute eye diameter; c, eye investment (for definitions, see methods). Eye investment was exponentiated in order that, in species which fall: (1) along the allometric fit, eye investment = 1x the investment predicted by the fit; (2) below the fit, eye investment < 1x; (3) above the fit, eye investment > 1x. Plot (d) above: Discrete traits (diel activity and turbidity; see key) shown by colours: darker colour, trait is present; lighter colour, trait is absent; half square, some species have trait, but not all. Generic or clade boundaries were adjusted according to the BEAST analysis results; note use of the terms ‘*xenophorid* group’, ‘*Lentigo* group’, ‘*Canarium* groups I-III’ and *‘Dolomena* group’ for ease of discussion. Plot (d) below: boxplot shows range of minimum (grey, right y axis) and maximum (black, left y axis) depths for stromboid genera or clade (where taxonomic boundaries are not formerly assigned). Upper and lower bounds of boxes are 25th and 75th percentiles; bars represent the median; outliers are black circles. Dashed lines at 200 m and 1,000 m delimit the dysphotic zone. Minimum and maximum depth were plotted at an identical scale and overlain using Adobe Illustrator. Number of species included in this study for each genus listed above plot.

Results from the *crunch* and *brunch* analyses suggested a significant association only between relative aperture width and maximum depth (Table 4). No significant association was found between diel activity and either group (small- or large-eyed clade) (Chi-square, χ*^2^* = 1.170, df = 1, p = 0.279) or genus (χ*^2^* = 30.385, df = 28, p = 0.345); however, significant associations were suggested between turbidity and both group (χ*^2^* = 22.61, df = 1, p<0.001) and genus (χ*^2^* = 55.00, df = 28, p = 0.02).

**Table 4.**
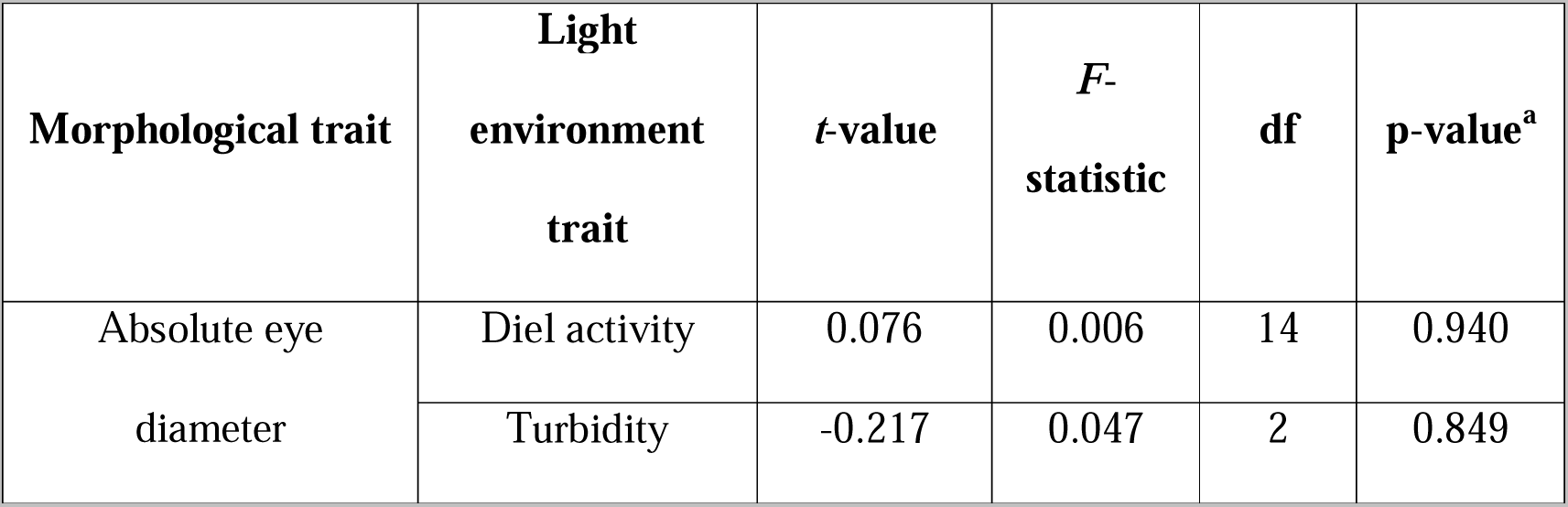
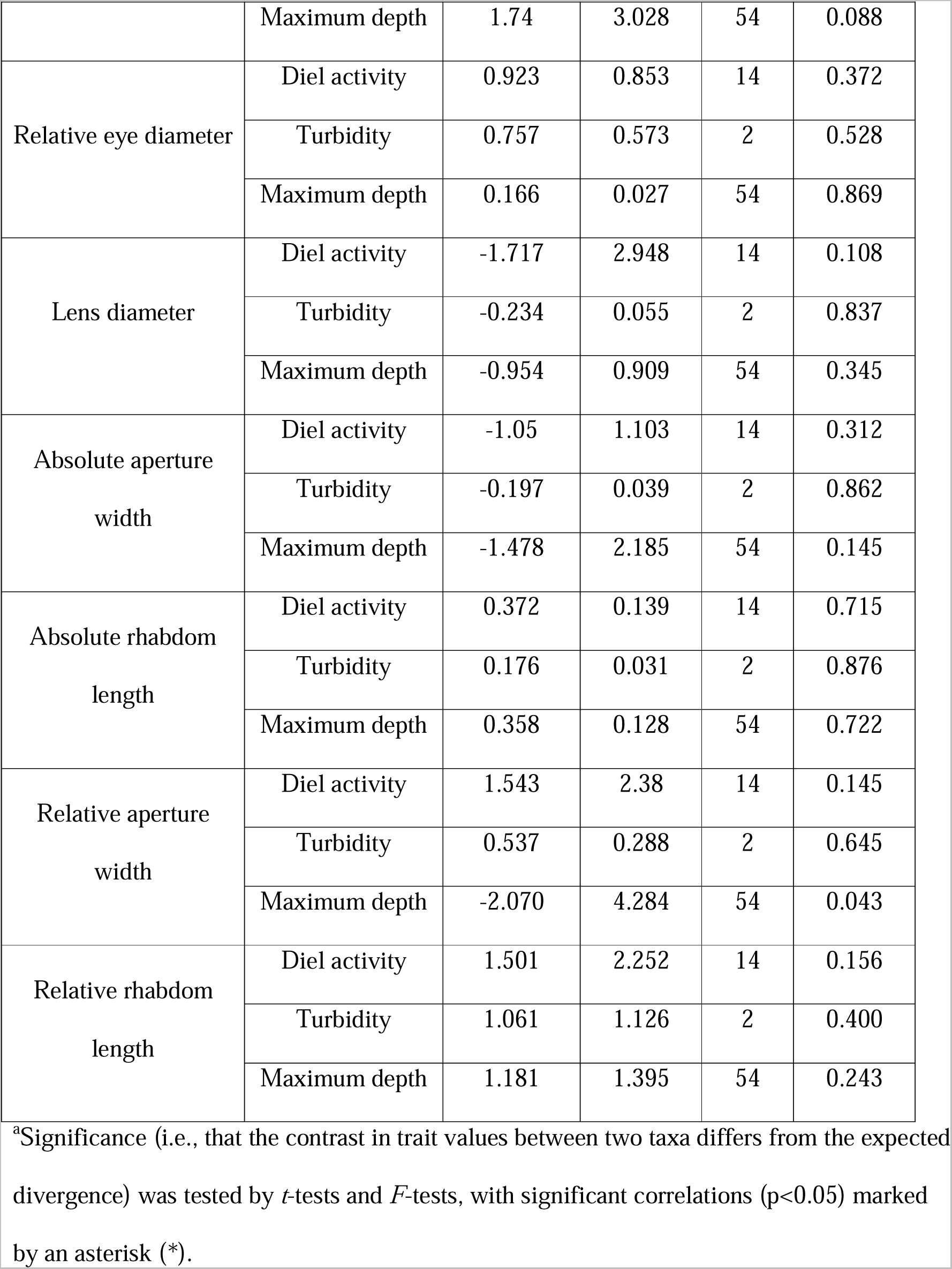
Phylogenetically Independent Contrast (PIC) Values Between Traits Relating to Light Environment and Log-transformed Morphological Traits.

| Morphological trait | Light environment trait | *t*-value | *F*-statistic | df | p-value<sup>a</sup> |
| --- | --- | --- | --- | --- | --- |
| Absolute eye diameter | Diel activity | 0.076 | 0.006 | 14 | 0.940 |
|  | Turbidity | -0.217 | 0.047 | 2 | 0.849 |
|  | Maximum depth | 1.74 | 3.028 | 54 | 0.088 |
| Relative eye diameter | Diel activity | 0.923 | 0.853 | 14 | 0.372 |
|  | Turbidity | 0.757 | 0.573 | 2 | 0.528 |
|  | Maximum depth | 0.166 | 0.027 | 54 | 0.869 |
| Lens diameter | Diel activity | -1.717 | 2.948 | 14 | 0.108 |
|  | Turbidity | -0.234 | 0.055 | 2 | 0.837 |
|  | Maximum depth | -0.954 | 0.909 | 54 | 0.345 |
| Absolute aperture width | Diel activity | -1.05 | 1.103 | 14 | 0.312 |
|  | Turbidity | -0.197 | 0.039 | 2 | 0.862 |
|  | Maximum depth | -1.478 | 2.185 | 54 | 0.145 |
| Absolute rhabdom length | Diel activity | 0.372 | 0.139 | 14 | 0.715 |
|  | Turbidity | 0.176 | 0.031 | 2 | 0.876 |
|  | Maximum depth | 0.358 | 0.128 | 54 | 0.722 |
| Relative aperture width | Diel activity | 1.543 | 2.38 | 14 | 0.145 |
|  | Turbidity | 0.537 | 0.288 | 2 | 0.645 |
|  | Maximum depth | -2.070 | 4.284 | 54 | 0.043 |
| Relative rhabdom length | Diel activity | 1.501 | 2.252 | 14 | 0.156 |
|  | Turbidity | 1.061 | 1.126 | 2 | 0.400 |
|  | Maximum depth | 1.181 | 1.395 | 54 | 0.243 |

<sup>a</sup>Significance (i.e., that the contrast in trait values between two taxa differs from the expected divergence) was tested by *t*-tests and *F*-tests, with significant correlations ( $p < 0.05$ ) marked by an asterisk (\*).

#### Large-eyed clade ((Rostellariidae + Seraphsidae) + Strombidae)

Independent *t*- tests suggested that the means of the following traits were significantly different between diurnal and cathemeral species in the large-eyed clade: body whorl width (*t* = 3.350, df = 41, p = 0.002), absolute eye diameter (*t* = 2.428, df = 41, p = 0.020), lens diameter (*t* = 2.836, df = 41, p = 0.007), relative rhabdom length (*t* = −2.787, df = 41, p = 0.008), relative aperture width (*t* = −3.425, df = 26.701, p = 0.002), and relative eye diameter (*t* = −2.672, df = 41, p = 0.011) (Fig. 4). *Terebellum* and *Varicospira* had a lower eye investment and smaller absolute eye diameter than the remainder of the large-eyed clade (Figs 1, 3b, c).

**Figure 4.**
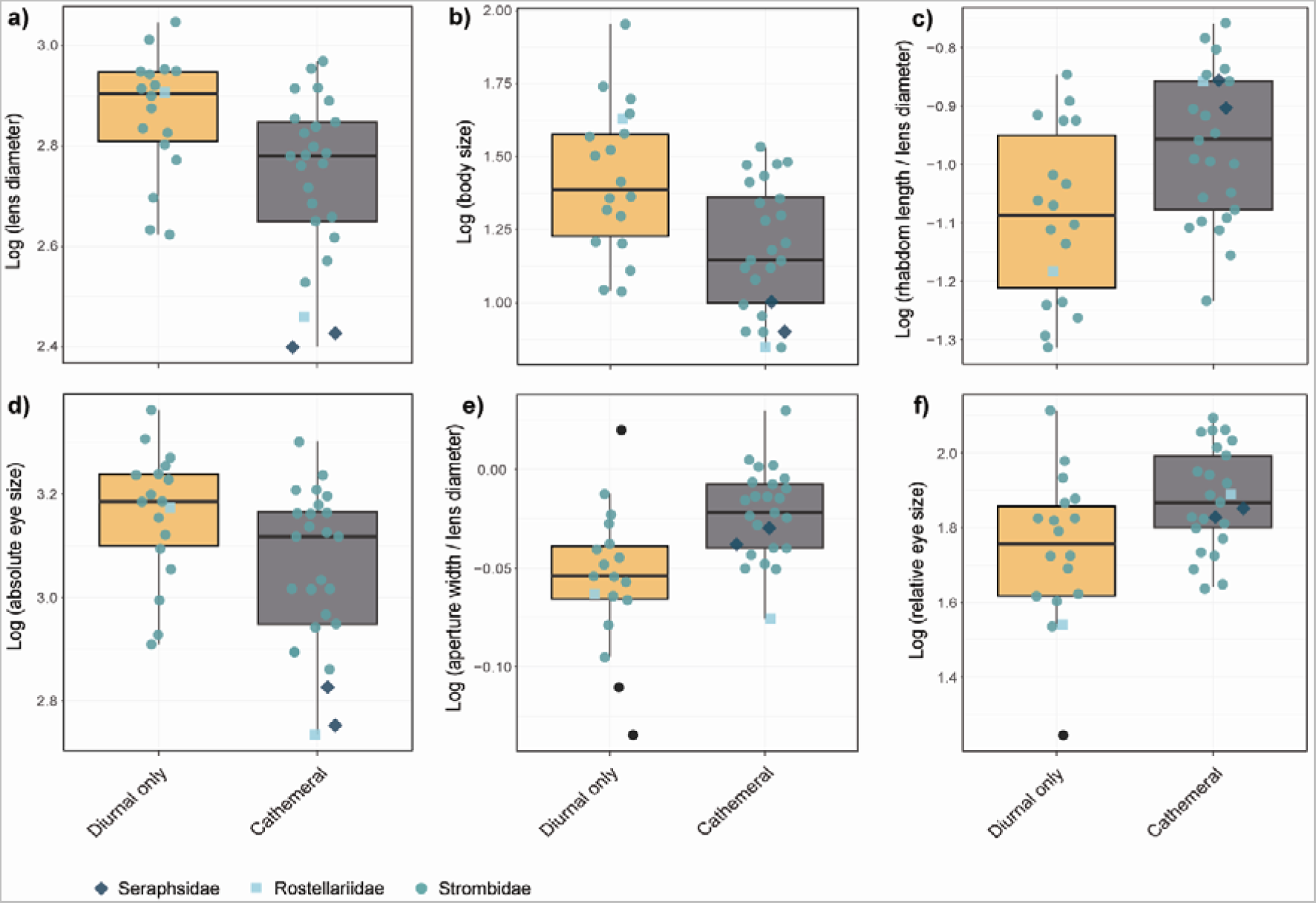
Variation in eye size and structure between (yellow) diurnal only and (grey) cathemeral species within large-eyed families Strombidae, Rostellariidae and Seraphsidae. Boxplots show: (a) lens diameter; (b) body size; (c) rhabdom length relative to lens diameter; (d) absolute eye diameter; (e) aperture width relative to lens diameter; (f) eye diameter relative to body size (see methods for full definitions of each trait). Scatter point shape and colour correspond to family (see key). The upper and lower bounds of the boxes are 25th and 75th percentiles, bars represent the median, and outliers are shown as black symbols, with shape also corresponding to family.

### Variation in Eye Structure across Stromboidea

Independent t-tests suggested that the means of the following traits were significantly different between small- and large-eyed clades: eye investment (*t*-test, *t* = 15.669, df = 54, p<0.001), absolute eye diameter (*t* = 9.960, df = 54, p<0.001), and relative eye diameter (t = 9.619, df = 54, p<0.001). On average, eye investment in the large-eyed clade was 1.76 ± sd 0.23 times larger (s.e.m 0.07, n = 48) than would be expected by body size, compared to mean 0.39 ± sd 0.21 times larger (s.e.m 0.03, n = 8) in the small-eyed clade. Among genera, there was a significant difference in absolute eye diameter (Kruskal-Wallis, *H* = 47.949, df = 28, p = 0.011) and relative eye diameter (*H* = 44.251, df = 28, p = 0.026), but narrowly not a significant difference for eye investment (*H* = 27.064, df = 28, p = 0.051). Tukey tests suggested significant differences among 23.2% of pairwise comparisons of eye investment among genera, with adjusted p values of <0.05, as well as 19.5% and 8.87% of comparisons of absolute and relative eye diameter, respectively.

ANCOVAs suggested that there was a significant difference in absolute eye diameter (*F* (2, 53) = 153.20, p<0.001) and relative eye diameter (*F* (2, 53) = 213.90, p<0.001) between the large- and small-eyed groups after controlling for body size (Fig. 5). There was also a significant difference in rhabdom length between groups controlling for lens width (*F* (2, 53) = 10.92, p<0.001) (Fig. 5). This suggests that absolute rhabdom length, and relative and absolute eye size, are proportionally different between the large- and small-eyed clades. This is evident in light microscopic comparisons, which also show a higher number of photoreceptors within the large-eyed clade (Fig. 6). Mean absolute rhabdom length was 60.6 ± sd 12.3 μm (s.e.m 1.8 μm, n = 8) in the large-eyed clade, compared to 21.0 ± sd 5.7 μm (s.e.m 2.0 μm, n = 8) in the small-eyed clade. Additionally, there was significant effect of lens width and group on aperture width, as well as a significant interaction between lens width and group (*F* (3,52) = 959.90, p<0.001), implying that the slope of the regression is different between the two clades (Fig. 5). The allometric slope for rhabdom length versus lens diameter was shallow in the large-eyed group, with a weak positive correlation (Pearson’s Correlation Coefficient, *r*= 0.359, *t* = 2.611, df = 46, p = 0.012) (Fig. 5). Observation of eyes using light microscopy revealed further differences between clades: lenses in the large-eyed clade were shattered and had a greater variation in staining across the lens (Fig. 6a–d), conversely, lens cores of small-eyed clade species were intact (Fig. 6e, f).

**Figure 5.**
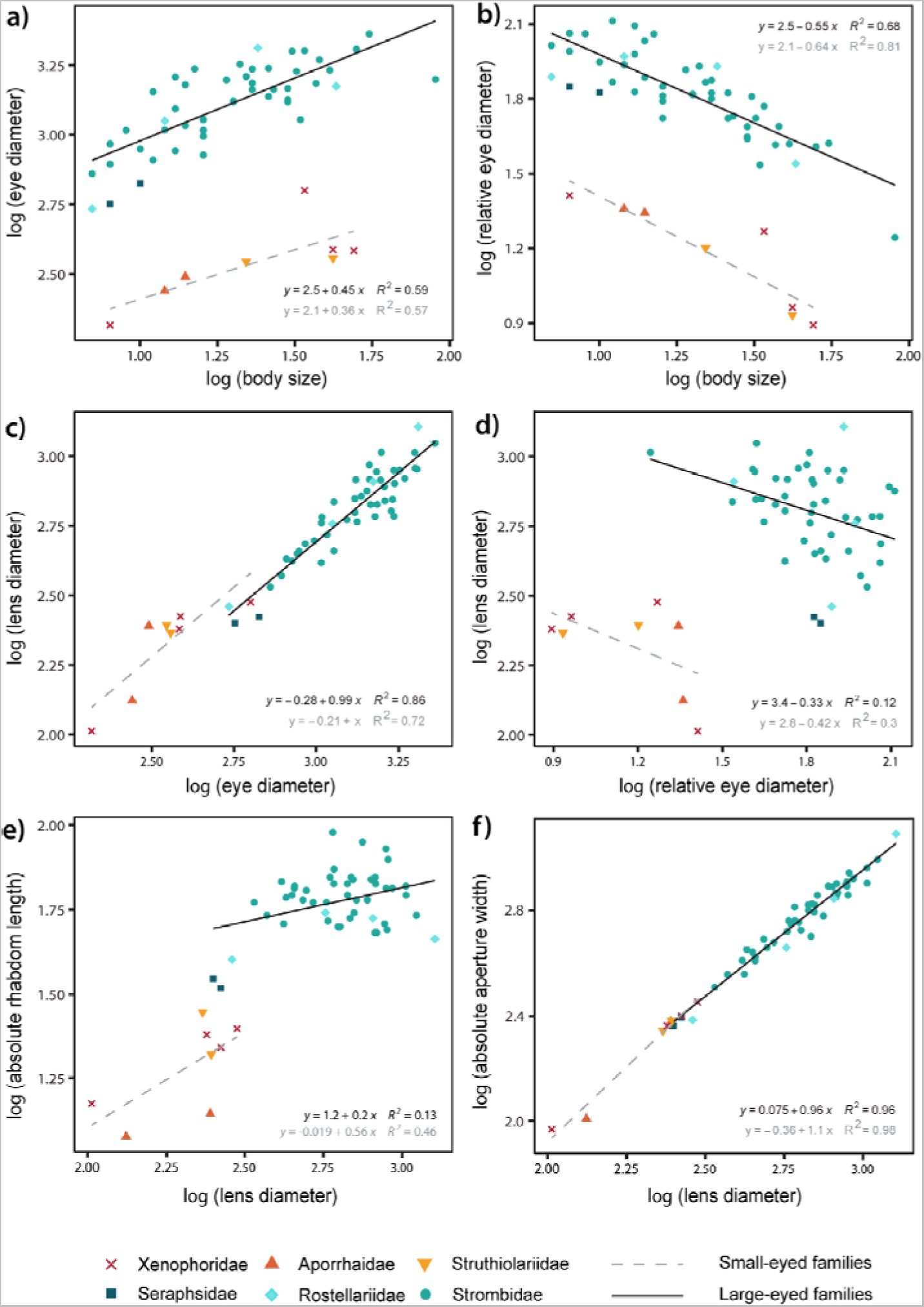
Scaling of eye size and structure, with linear models plotted for small- (grey dashed line) and large-eyed (black solid line) families. Panels show log-transformed data for: a, eye diameter relative to body size; b, relative eye diameter relative to body size; c, lens diameter relative to eye diameter; d, lens diameter relative to relative eye diameter; e, rhabdom length relative to lens diameter; f, aperture width relative to lens diameter (see methods for full definitions of each trait). Scatter point shape and colour correspond to family (see key).

**Figure 6.**
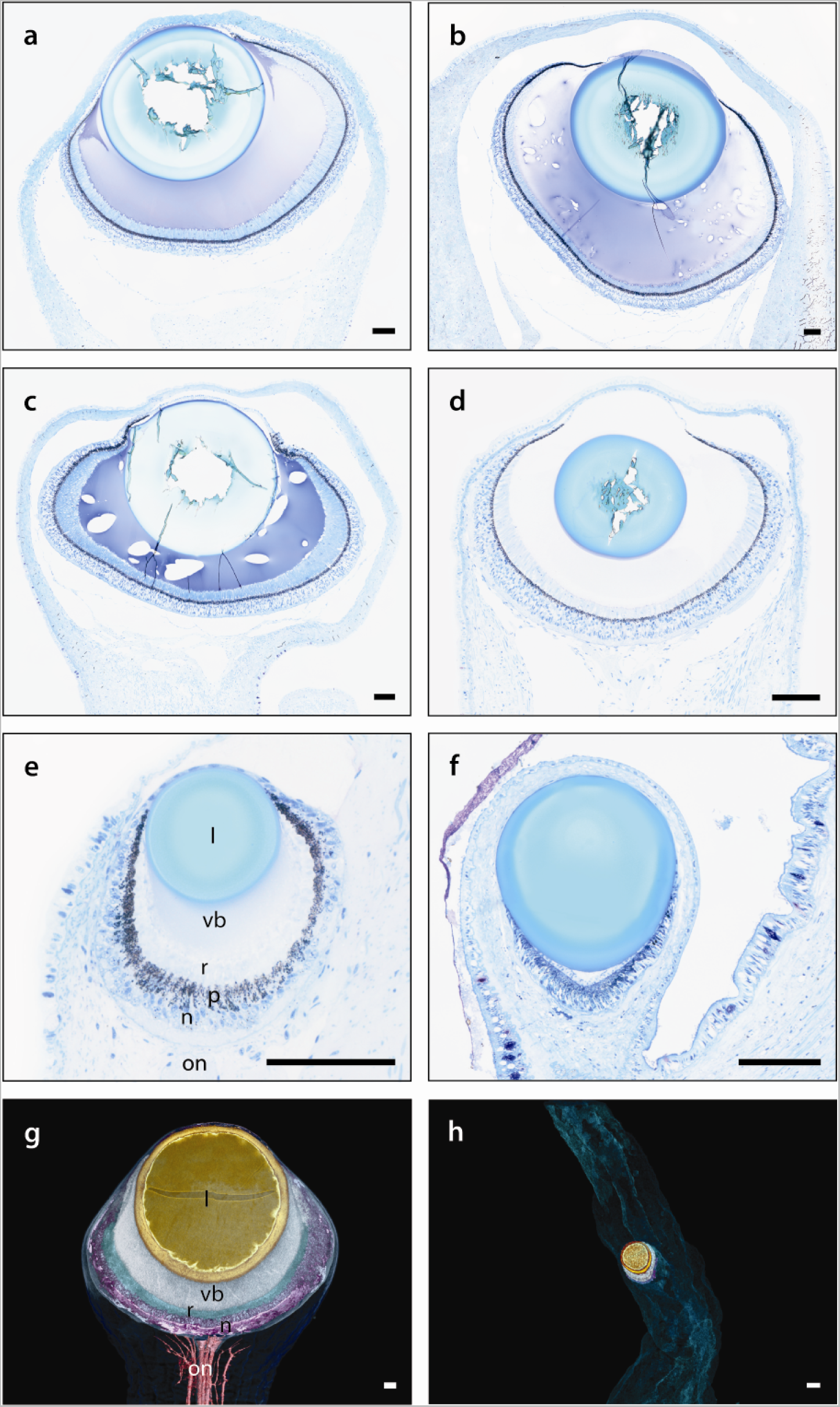
(a-f) Light micrographs showing histological sections through the centre of eyes and (g, h) μCT slices through the centre of tomographic models of eyes for the following species: a, strombid *Ministrombus variabilis*; b, strombid *Harpago chiragra*; c, strombid *Tridentarius dentatus*; d, seraphsid *Terebellum terebellum*; e, xenophorid *Xenophora solarioides*; f, h, aporrhaid *Aporrhais pespelecani*; g, rostellariid *Rostellariella delicatula.* Images (g, h) show differences in position of eyes on eyestalks, as well as in eye size (identical scale given). Abbreviations (shown in light micrograph image (e) and μCT image (g)): l, lens; n, nuclear layer; on, optic nerve; p, pigmented region; r, rhabdomeric layer; vb, vitreous body). Supplementary Figure S3 (available on Dryad) for μCT scan videos. Scale bar: 100 μm.

## Discussion

### Evolution of Visual Systems in Stromboidea

The comparatively large eyes of strombids, rostellariids and seraphsids stand out among gastropods which tend to have small eyes and low acuity vision (e.g., Hamilton and Winter 1984; Seyer, 1992, 1998; Gál et al. 2004). Due to lack of specimen availability, the largest strombids, the aptly-named *Aliger gigas* and *Titanostrombus goliath*, could not be included in this study. Even so, the largest eye diameter measured was 2.3 mm (strombid *Thersistrombus thersites*), almost twice the eye size of the predatory pterotracheoid gastropod *Oxygyrus keraudreni* (1.2 mm; Land, 1982). Although the latter likely has a larger eye size relative to its body size, this is still a surprising comparison given the non-predatory lifestyle of the strombids. In this study, ancestral state reconstruction suggested that the evolution of large eyes in Stromboidea was followed by a further increase in absolute eye size within two separate clades of the large-eyed group (Fig. 1). Limited sample sizes from some groups mean that reversal cannot be excluded as a possibility; however, parallel evolution is supported by differences in scaling between the Seraphsidae and *Varicospira*, and the larger-eyed strombids and remainder of rostellariids (Figure 5d, e). The evolutionary drivers of these large eyes will be discussed below, bearing in mind the trade-off between the energetic costs these large eyes incur and the competitive advantages they must surely provide with respect to resolution and sensitivity (Marshall, 2017).

Eyes within the large-eyed clade likely provide much finer resolution than those of the small-eyed clade, due to longer focal lengths and higher numbers of photoreceptor cells (Fig. 6). Furthermore, morphological studies suggest a graded refractive index in the lenses of the large-eyed group, which would reduce spherical aberration (Seyer 1994; Land and Nilsson 2012). Light microscopy reveals lenses with highly inhomogeneous staining in all large-eyed species, and shattered cores as would be expected in very hard tissues (Seyer 1994; Caropreso et al. 2000; Fig. 6). By contrast, the lens cores within small-eyed specimens sectioned remained intact, suggesting a more homogenous lens structure prone to spherical aberration (Fig. 6) (Seyer 1994; Blumer 1996; Land and Nilsson 2012). High visual acuity in strombids is supported by both behavioural and anatomical studies of strombid *Conomurex luhuanus* (Seyer et al. 1994; Irwin et al. 2022); however, no behavioural experiments have yet investigated differences in visual capabilities between the large- and small-eyed groups. Nevertheless, the spatial resolution of *C. luhuanus*, approximately 1°, is almost four times finer than that of the littorinid *Littorina littorea* (Seyer et al. 1992, 1994; Irwin et al. 2022). The latter has an eye size of ca. 240 μm, of a similar magnitude to that of the small-eyed stromboid clade (mean eye size: 363 ± 124 μm). Therefore, we can predict that the eyes of struthiolariids, xenophorids and aporrhaids may provide similarly lower acuity vision (Seyer et al. 1992) (Fig. 1; Supplementary Table S3).

A higher visual sensitivity in the large-eyed stromboid clade is supported by their eyes having wider apertures and longer rhabdom lengths (Fig. 6; Supplementary Table S3). These longer rhabdoms (mean 60.6 ± sd 12.3 μm, compared to 21.0 ± sd 5.7 μm in the small-eyed clade) extend the path of light travelling through the photoreceptors, increasing photon absorbance (Land and Nilsson 2012). There is also a difference in rhabdom structure between clades; in the large-eyed clade, rhabdoms are comprised of a cytoplasmic core surrounded by circularly arranged microvilli (Gillary and Gillary 1979; Irwin et al. 2022). This structure is a key adaptation for longer absolute rhabdom length, as this allows more efficient transport to the microvilli and provides structural support, aided by the long filaments of sensory cells (Hughes 1976). By contrast, distal processes in the small-eyed clade are comprised of microvilli extending straight towards the lens from an apical surface (Blumer 1996); similar structures in other gastropods are not known to exceed 25–35 μm in length which is consistent with our results (Supplementary Table S3) (Hughes 1970). Furthermore, the microvillar orientation in small-eyed groups (parallel to incident light passing through the centre of the lens, as opposed to perpendicular in the large-eyed clade) likely supports less efficient photoreception, reducing sensitivity (Hughes 1976; Blumer 1996).

### Adaptations to Different Photic Environments

Depth is a key determinant of light availability in the marine environment (Warrant 2004). This is reflected in our results, which provide evidence of a significant correlation between maximum depth and relative aperture width within Stromboidea (Table 4). A wide aperture relative to focal length increases sensitivity at the cost of resolution; therefore, this is likely an adaptation to lower light availability in deeper waters. By contrast, a larger absolute eye size may be an adaptation for fine resolution in brighter environments, given that the large-eyed stromboid group generally inhabits shallower maximum depths (Figs 2a, 3). However, a divergent strategy may be illustrated by the large eyes of rostellariids *Rimellopsis* and *Rostellariella* and strombid *Dolomena labiosa*, which inhabit deeper waters. In such lower light environments, natural selection can either reduce investment in a visual system, even abandoning the system entirely, or increase investment to improve sensitivity (Howland et al., 2004; Dugas and Franssen, 2012). No significant correlations were found between turbidity and eye size; however, this may be due to the lower taxon sampling within the small-eyed clade, and the fact that turbidity data were collected from satellite data and not from collection sites. Nevertheless, it is worth noting that small-eyed stromboid taxa are all light-limited, either due to high turbidity, which reduces light availability even in shallow waters (Warrant 2004), to a deeper depth range, or to both (Figs 1, 3).

Within the large-eyed stromboid clade, a larger lens diameter relative to both aperture width and rhabdom length are present in species that are only active during the day, such as *Harpago chiragra* (Figs 4, 6c, e). These morphological traits are associated with higher acuity due to a longer focal length, and therefore prioritising sensitivity over resolution (Hall and Ross 2007; Land and Nilsson 2012). By contrast, wider apertures and longer rhabdoms relative to lens size are present in cathemeral groups, such as *Ministrombus variabilis* and *Tridentarius dentatus* (Figs 4, 6a, c). These traits increase the number of photons absorbed, adaptations that are well-described in nocturnal insects, and suggest prioritising sensitivity over resolution (Land et al. 1999; Greiner et al. 2004; Frederiksen and Warrant 2008). These cathemeral animals also possess a smaller body size than those only active during the day (Fig. 4); a smaller shell, and therefore presumably thinner and mechanically weaker, is more vulnerable to visually guided shell-crushing and shell-peeling predators such as cephalopods, crustaceans, fish and turtles (Supplementary Table S6 available on Dryad) (e.g., Randall, 1964; Stoner, 1994; Delgado, 2002; Üstüner, 2018). Therefore, cathemeral rather than diurnal behaviour may limit exposure to the increased predation pressure during the day.

### Predation Pressure as a Driver of Visual Evolution

A greater investment in visual acuity and sensitivity requires more extensive visual processing, with higher energetic costs (Laughlin et al. 1998; Meyer-Rochow 2001; Niven and Laughlin 2008). Large-eyed stromboids had a significantly larger eye investment, with eyes that were almost twice the expected value for their body size (based on Stromboidea as a whole), compared to the small-eyed clade which had eyes less than half the expected size. These larger, more metabolically expensive eyes likely only evolved in stromboid groups that benefitted from an increased quality of visual information, such as those subject to predation pressure, as has been suggested for several other species with visual predators (including some birds [Møller and Erritzoe 2010], crustaceans [Glazier and Deptola 2011] and squid [Nilsson et al. 2012]). In gastropods, a strong selection pressure from predation may result in a variety of metabolically expensive defensive adaptations, either by increasing armour as a passive defence, or by facilitating an active escape (Vermeij 1977, 1987; Harper 2022). An increase in the variety and number of defensive traits during the Mesozoic is often linked with a rise in durophagous predators, one of many changes to the marine benthic community which characterised the Mesozoic Marine Revolution (Vermeij 1977, 1987). This includes shell traits found within stromboids (thickened shell lip, ornamentation, and/or camouflage), which Roy (1994) suggested are associated with increased predation pressure. The increase in absolute eye size within Stromboidea is also estimated to have occured during the Mesozoic (Fig. 1), although at present no data exist to link this to the rise of specific durophagous taxa. The contrast in predation pressure between the large- and small-eyed groups may be associated with differences in depth and latitude, as is suggested for other prey taxa (e.g., Harper and Peck 2016; Ashton et al. 2022). Generally, there is assumed to be a higher predation pressure on tropical gastropods, indicated by increased breakage-induced shell repair and a higher turbinid operculum thickness in warmer waters than in temperate regions (Vermeij et al. 1981; Vermeij 1987; Alexander and Dietl 2003; Vermeij and Williams 2007; Stafford et al. 2015). As the majority of large-eyed stromboid species inhabit shallow tropical regions, these are likely to encounter more predators than the smaller-eyed species, which tend to inhabit deeper and/or temperate waters (Morton 1951; Ponder 1983; Roy 1996; Briggs 1999).

Another key difference between the large- and small-eyed stromboid families is the placement of the eyes. Within large-eyed families, eyes are located at the ends of long and mobile ommatophores. This characteristic is uncommon in the caenogastropods, otherwise known only in the predatory Conidae and Terebridae (Simone 2011; Ponder and Lindberg 2020). By contrast, the much shorter ommatophores of the small-eyed families are limited in movement, with the shell obscuring most of the visual field (Simone 2005). A larger visual field means that more information can be extracted from an animal’s surroundings, an important adaptation for visually guided behaviour such as the predator avoidance behaviour noted in strombids (Martin et al. 2007; Martin 2014; Lisney et al. 2020; Irwin et al., 2021). Predation pressure therefore may be a potential driver for the selection of these long, mobile eyestalks. Similar strategies are described with respect to the long, mobile eyestalks of crabs and the dorsally positioned eyes of benthic fishes (Land and Layne 1995; Zeil and Hemmi 2006; Bellwood et al. 2014; Lisney et al. 2020). In this study, ancestral state reconstruction resolved the root node state for the location of eyes along tentacles in Stromboidea as most likely being at the base of the tentacle, albeit with low support (p = 0.64), suggesting that the position of eyes on tentacles may have co-evolved with increased absolute eye size (Fig. 1).

In strombids, withdrawal into the shell can be initiated in behavioural studies by rapidly expanding visual stimuli that mimic fast-approaching potential predators (Irwin et al. 2022). However, strombids alone are known to also possess a second avoidance strategy in the ability to perform a series of fast, directional backward leaps away from predatory cone snails (Kohn and Waters 1966); limited information is available on similar rapid locomotion in seraphsids (Savazzi 1991) (Supplementary Table S1). This leaping escape response is presumably effective, given that *Lambis lambis* can reach speeds of 11.9 mm/s in comparison to the crawling rate of 0.2 mm/s for the predatory cone snail *Conus agassizii.* Experiments using cone snail extract found that the escape response can be triggered by chemosensory stimuli only; however, it is not clear from other experiments (wherein a strombid was placed near to a cone snail) whether this response can be also triggered exclusively by visual stimuli (Kohn and Waters 1966; Field 1977; Watson et al. 2014; Lefevre et al. 2017). Nevertheless, one such behavioural study suggested that vision is at least important in maintaining the direction of the escape response (Field 1977). When eyes were removed, strombids leapt away from the predator down a chemical gradient, but eventually arced back towards the cone snail, thus rendering the escape response ineffective (Field 1977).

Within Stromboidea, all families possess an operculum on the distal end of the foot, extending at least partly beyond it, which facilitates the jerky, discontinuous leaps that are characteristic of normal stromboid locomotion (Simone 2005). Nevertheless, differences in operculum shape and foot morphology between the small- and large-eyed clade may amount to contrasts in predator avoidance responses. Only strombid and xenophorid behaviours have been directly compared so far, wherein Berg (1975) noted that, unlike strombids, *Xenophora conchyliophora* does not produce a rapid escape response and only withdraws into the shell when threatened. There is also a difference in righting behaviour; when overturned, strombids can right the shell with a rapid kick of the foot, while *X. conchyliophora* uses a slow, gradual motion, more typical of other gastropods (Berg 1975). This could be due in part to an overall difference in shell shape (fusiform versus trochiform; Simone 2005); however, variation in locomotion may also be attributed to differences in foot and operculum morphology. Strombids and seraphsids possess a long foot lacking a distinct crawling sole, with an operculum that is elongated, pointed, and often highly serrated (rostellariids were not included in this comparative study, but possess a similar operculum; Simone, 2005). By contrast, small-eyed families possess a comparatively small and stubby foot with a crawling sole, as well as an operculum that is more elliptical in shape (Simone 2005). Therefore, variation in foot morphology, absolute eye size, and locomotory capabilities may reflect possible co-evolution between visual and locomotory systems.

## Conclusions

Molluscan eyes vary widely across the phylum in both size and structure, with a range of adaptations to the surrounding environment. As measured so far, strombids have larger eyes than any other gastropod, reaching up to 2.3 mm in diameter according to this study. Ancestral state reconstruction indicates that a further increase in eye size above 1 mm occurred independently within two separate stromboid clades. This study also suggests that the light environment as determined by maximum depth is a key factor in the evolution of relative aperture width within this superfamily. Large apertures relative to lens size, indicating increased sensitivity, are associated with eyes from species inhabiting deeper maximum depths with lower light availabilities. Additionally, our results find that eye structure varies significantly with differences in diel activity within the large-eyed stromboid clade. Larger lenses relative to aperture widths and rhabdom lengths, reflecting a higher investment in resolution over sensitivity, are more common in animals active only during the day. These diurnal animals are also larger in body size; those with a smaller body size are more likely to be cathemeral, possibly because a smaller and presumably thinner shell increases vulnerability to shell-crushing and shell-peeling predators.

Overall, this study suggests that evolution of larger eyes and higher visual performance was at least partly driven by key visual tasks: early detection and avoidance of predators. Owing to the rapid escape response known to occur only in large-eyed taxa, and variations in foot morphology between large- and small-eyed groups, larger eyes and improved visual performance is suggested to have co-evolved with rapid locomotion. This is a new finding implicating predation pressure as a potential driver in the evolution of visual and locomotory systems within a gastropod group.

## Acknowledgements

Part of the material used in this study originates from several research cruises and expeditions organized by the MNHN and ProNatura International as part of the *Our Planet Reviewed* program, and by the MNHN and the Institut de Recherche pour le Développement as part of the *Tropical Deep-Sea Benthos* program (KAVIENG 2014 (dx.doi.org/10.17600/14004400), and PAPUA NIUGINI (dx.doi.org/10.17600/18000841) in Papua New Guinea; CONCALIS (dx.doi.org/10.17600/8100010), EXBODI (dx.doi.org/10.17600/11100080) and KOUMAC 2.3 in New Caledonia; Dakar’09 in Senegal; ATIMO VATAE (dx.doi.org/10.17600/10110040) and MIRIKY in Madagascar; INHACA 2011 in Mozambique; MADIBENTHOS in Martinique – more information can be found at expeditions.mnhn.fr). These expeditions operated under the regulations then in force in the countries in question and satisfy the conditions set by the Nagoya Protocol for access to genetic resources. The authors thank Philippe Bouchet and Nicolas Puillandre for access to this material, as well as Virginie Héros, Barbara Buge and Julien Brisset (MNHN) for their help in curating the vouchers. We also thank Nicolas Puillandre for the additional COI sequences included in this study (see Supplementary Material 1). We thank Andreia Salvador for the access of NHMUK specimens and in help creating registration numbers. For the loan of museum specimens, we also thank John Slapcinsky (University of Florida), Richard Willan (Museum and Art Gallery of the Northern Territory, Australia), Serge Gofas (University of Málaga), Tan Koh Siang and Tan Siong Kiat (National University of Singapore). We thank Gijs Kronenberg (Naturalis Biodiversity Center, Netherlands) for confirming species identification and discussion of key morphological characters for stromboid groups. We are grateful to Elena Lugli and Nathan Kenny for initial help in the lab, and Claire Griffin of the NHMUK Sequencing Facility for completing the PCR clean-up and Sanger sequencing. We thank Vincent Fernandez and Brett Clarke (NHMUK) for performing μCT scans and providing access to VGStudio Max software. Finally, we are grateful to Freya Goetz (National Museum of Natural History, NMNH) for training on histological techniques.

## Data Availability

All Supplementary Data are available in the Dryad Digital Repository, at: https://datadryad.org/stash/share/Nm-w5jOQWmN1Ny5puAKns7i6Lvf_ghcxslDSPD1y29s [doi:10.5061/dryad.pnvx0k6v0].

